# From Cardiac Myosin to the Beta Receptor: Autoantibodies Promote a Fibrotic Transcriptome and Reduced Ventricular Recovery in Human Myocarditis

**DOI:** 10.1101/2024.06.19.599804

**Authors:** Jennifer M. Myers, Clayton Sandel, Kathy Alvarez, Lori Garman, Graham Wiley, Courtney Montgomery, Patrick Gaffney, Stavros Stavrakis, DeLisa Fairweather, Katelyn A. Bruno, Yan Daniel Zhao, Leslie T. Cooper, Madeleine W. Cunningham

**Author notes:** These authors contributed equally to this work and share first authorship (Jennifer M. Myers, Clayton Sandel). Address for Correspondence: Madeleine W. Cunningham, PhD, University of Oklahoma Health Sciences Center, 975 NE 10th Street, Biomedical Research Center Room 217, Oklahoma City, OK 73104.

## Abstract

**Background:** Myocarditis leads to dilated cardiomyopathy (DCM) with one-third failing to recover normal ejection fraction (EF50%), and there is a critical need for prognostic biomarkers to assess risk of nonrecovery. Cardiac myosin (CM) autoantibodies (AAbs) cross-reactive with the β−adrenergic receptor (βAR) are associated with myocarditis/DCM, but their potential for prognosis and functional relevance is not fully understood.

**Methods:** CM AAbs and myocarditis-derived human monoclonal antibodies (mAbs) were investigated to define pathogenic mechanisms and CM epitopes of nonrecovery. Myocarditis patients who do not recover ejection fraction (EF<50%) by one year were studied in a longitudinal (n=41) cohort. Sera IgG and human mAbs were investigated for autoreactivity with CM and CM peptides by ELISA, protein kinase A (PKA) activation, and transcriptomic analysis in H9c2 heart cell line.

**Results:** CM AAbs were significantly elevated in nonrecovered compared to recovered patients and correlated with reduced EF (<50%). CM epitopes specific to nonrecovery were identified. Transcriptomic analysis revealed serum IgG and mAb 2C.4 induced fibrosis/apoptosis pathways *in vitro* similar to isoproterenol treated cells. Sera IgG and 2C.4 activated PKA in an IgG and βAR-dependent manner. Endomyocardial biopsies from myocarditis/DCM revealed IgG+ trichrome+ tissues.

**Conclusions:** CM AAbs were significantly elevated in nonrecovered patients, suggesting novel prognostic relevance. CM AAbs correlated with lower EF, and Ab-induced fibrosis/apoptosis pathways suggested a role for CM AAbs in patients who do not recover and develop irreversible heart failure. Homology between CM and βARs supports mechanisms related to cross-reactivity of CM AAbs with the βAR, a potential AAb target in nonrecovery.

## Introduction

Myocarditis is a rare inflammatory sequela of viral infections that progresses to dilated cardiomyopathy (DCM), heart failure (HF), transplantation, and death in one-third of patients (1–3), and contributes to a substantial number of sudden deaths in young (< 40 years) adults (4). Definitive diagnosis of myocarditis is performed by the Dallas Criteria during histopathological examination of endomyocardial biopsy (EMB) (5), however sampling error reduces the sensitivity. Prognosis of myocarditis patients is a critical need, with few biomarkers that predict outcomes early in disease (3). Heart-directed autoantibodies (AAbs) have been associated with greater cardiac risk but the mechanisms of autoantibody-mediated heart injury remain unknown (6–11).

Multiple studies indicate AAbs may be critical in the pathogenesis of myocarditis. The presence of cardiac-directed AAbs was predictive of outcomes in an immunosuppressive therapeutic trial with retrospective analysis of myocarditis patients (12), and purified anti-CM IgG induced myocarditis with inflammatory cell infiltrate as well as DCM when passively transferred into BALB/c mice or Lewis rats. (13,14). Although several different specificities of AAbs have been described by Maisch (15), Kaya (16), and others (17–19), none are currently used as biomarkers for diagnosis or prognosis of disease. CM AAbs have been associated with DCM, fibrosis, and HF (2,20–26), and CM is one of the few heart proteins which can induce myocarditis when administered to susceptible animal models (7,27–32). A recent study suggests that elevated serum AAbs against the heart, female gender, fulminant onset, and lower ejection fraction are all predictors of death or transplantation in myocarditis before the introduction of immunosuppressive therapies(33).

Our previous studies of experimental autoimmune myocarditis (EAM) in Lewis rats demonstrated *in vivo* IgG myocardial deposition as a hallmark of myocarditis pathology (14). *In vivo* IgG deposition in heart tissue also followed passive transfer of serum IgG purified from rats immunized with CM. IgG deposition and dilated cardiomyopathy in heart tissues has also been reported in mouse models of CM-induced myocarditis (34), but not linked to poor outcomes or heart function. Mouse models of viral myocarditis (MCMV and CVB3) have also reported CM AAb in the serum during acute and chronic myocarditis as well as cardiac deposition of CM AAb (35,36). Mouse models demonstrated IL-17A as an essential cytokine for the progression of myocarditis to DCM (37), and in orchestrating fibrosis in myocarditis and other diseases (38). IL17A was suggested as a biomarker elevated in the nonrecovered phenotype in human myocarditis where Th17 responses were linked to heart failure(39). CM AAbs have also been associated with Th17 responses in other types of inflammatory heart diseases (40–45) including poor outcomes in type 1 diabetes (46,47).

CM AAbs were shown by our laboratory to cross-react with β-adrenergic receptors (βAR), and passive transfer of CM AAbs caused cardiomyocyte apoptosis and DCM in the Lewis rat model of myocarditis, resulting in cAMP-dependent protein kinase A (PKA) signaling activation in cardiomyocytes (14,20). CM AAbs in the Lewis rat model were specific for βAR as they did not react with α-adrenergic receptors (αARs) (14). Similarly, Jahns et al demonstrated pathogenicity of AAbs against the β-adrenergic receptors (βAR) where passive transfer of sera induced dilated cardiomyopathy in a mouse model of EAM (18). Other studies have shown anti-βAR AAbs associated with idiopathic arrhythmias and myocardial sites of focal infiltration and necrosis in human biopsy tissue (48). β-blocker therapy with carvedilol was efficacious in chronic heart failure patients with AAbs to the β-adrenergic receptor (βAR) (49).

While approximately 70% of patients with myocarditis recover left ventricular ejection fraction (LVEF), biomarkers to identify patients who will not spontaneously recover their LVEF is an unmet clinical need to inform treatment decisions and prevent progression to DCM, HF, and transplant (50). Our present study provides evidence suggesting CM AAbs may be prognostic for poor outcomes in myocarditis/DCM patients via βAR activation, and that CM AAbs cause transcriptional changes that may promote cardiac fibrosis and apoptosis. A human monoclonal antibody (mAb) and sera derived from myocarditis/DCM patients activated βAR and induced elevated expression of fibrosis and apoptosis genes. We show that CM AAbs are elevated at clinical presentation in patients who fail to recover LVEF by one year, and negatively correlate with LVEF in patients with poor EF outcomes. IL-17A, a serum biomarker associated with the nonrecovered human myocarditis phenotype, was previously correlated with poor outcomes (6) and was detected in endomyocardial biopsies and correlated with IgG deposition and concomitant fibrosis. In total, we provide further evidence that supports the hypothesis that CM AAbs contribute to pathogenesis of human myocarditis and may be important in identifying myocarditis patients who fail to recover normal heart function.

## Methods

### Patient Cohort and Experimental Design

Forty-one adults (age 18–89 years, 66% male) with a diagnosis of acute myocarditis/DCM and 32 healthy adult volunteers (age 18–69 years, 59% male) were enrolled in the study. Timing of blood collection, deaths, or dropouts resulted in sample sizes of less than 41 patients in some assays. All myocarditis/DCM patients were enrolled less than 6 months from date of symptom onset, met clinical criteria for myocarditis, had ejection fractions less than 50%, and had no greater than 50% stenosis in any epicardial coronary artery (1). Patients were further characterized by (a) endomyocardial biopsy that met Dallas criteria for myocarditis/borderline myocarditis, (b) cardiac MRI scan that met consensus conference diagnostic criteria for acute myocarditis, or (c) echocardiogram demonstrating LVEF less than 50% with an otherwise unexplained rise in troponin. All subjects had clinical evaluation and blood collection during their first patient visit (baseline), and blood from a subset of patients was collected at 1 to 3 and 6 to 12 months after baseline. Serum was stored at –80°C. Viral myocarditis was not determined in the cohort by PCR or viral culture from peripheral blood or heart biopsies.

### Antibody Staining of Heart Biopsies

Goat anti-human IgG (10 μg/ml), or isotype-control human IgG Ab (10 μg/ml; Sigma-Aldrich) was incubated on de-paraffinized tissue sections overnight at 4^°^C. Biotin-conjugated rat anti-goat IgG Ab (1/500; Jackson ImmunoResearch Laboratories) was incubated on tissues for 30 min. Alkaline phosphatase-conjugated streptavidin (Jackson ImmunoResearch Laboratories) was incubated on tissues at 1 μg/ml for 30 min at room temperature. Ab binding was detected with Fast Red substrate (BioGenex) against a counterstain of Mayer’s hematoxylin (BioGenex). Cell surface-bound Abs were detected with specific biotin-conjugated secondary Ab (1/500; Sigma-Aldrich).

### Trichrome Staining of Heart Biopsies

Tissue sections were stained in our laboratory using Masson’s Trichrome Method. Briefly, tissues were de-paraffinized and rehydrated through alcohol gradients, washed in distilled water, then re-fixed in Bouin’s solution [saturated picric acid, formaldehyde (37-40%), glacial acetic acid] for 1 hour at 56° C, or overnight at room temperature when formalin fixed. The tissues were cooled and washed in running water to remove yellow color, then rinsed in distilled water. Sections were then stained in Wiegert’s iron hematoxylin solution [stock solution A: hematoxylin and 95% alcohol; stock solution B: 29% ferric chloride in water, distilled water, concentrated hydrochloric acid; working solution -equal parts of stock solutions A and B for 10 minutes, washed in warm running water for 10 minutes, and rinsed in distilled water. The tissues were then stained in Biebrich scarlet-acid fuchsin solution (Biebrich scarlet, 1% aqueous, acid fuchsin, 1% aqueous, and glacial acetic acid) for 10 minutes, then rinsed in distilled water. Tissues were differentiated in phosphomolybdic-phosphotungstic acid solution (5% phosphomolybdic acid, 5% phosphotungstic acid) for 10 minutes. The solution was discarded, and tissue sections were stained with aniline blue solution (aniline blue, glacial acetic acid, distilled water) for 5 minutes. Sections were briefly rinsed in distilled water, placed in 1% glacial acetic acid solution for 3 minutes, washed in distilled water, then dehydrated in 95% ethyl alcohol, absolute ethyl alcohol, and cleared in xylene, two changes each. Sections were mounted with Permount. In trichrome-stained tissue sections, nuclei stain black and collagen stains blue. Cytoplasm, keratin, muscle fibers and intercellular fibers stain red.

### ELISA

Antigen human cardiac myosin (CM) prepared as previously described (41). Peptides of the S2-fragment were diluted at 10 µg/mL in carbonate/bicarbonate buffer and 50µL/well placed into each well of a 96-well ELISA plate. The plates were covered in plastic wrap and incubated overnight at 4^0^C. The plates were washed 5X with PBS/0.5% Tween. The plates were blocked by dispensing 100µL/well of 1% BSA/PBS and incubating for one hour at 37^0^C. The plates were washed as described above and 50µL/well of diluted (1:100) sera placed on the plate in duplicate wells. The plates were washed as described above and 50µL/well of diluted (1:100) sera placed on the plate in duplicate wells. The plates were covered in plastic wrap and placed overnight at 4^0^C. The following day, the plates were washed. Secondary antibody conjugated to alkaline phosphatase was diluted to 1:500 in 1% BSA/PBS and 50µL/well added and plates left to incubate for one hour at 37^0^C. The plates were then washed, and substrate solution added at 50µL/well. The optical density was read at an absorbance of 405nm after 15, 30, 60, and 90 minutes. Results shown in graphs are from the 60min timepoint.

### S2 Peptide Synthesis

Peptides spanning the S2 fragment (hinge region) of the human cardiac myosin molecule were synthesized and purified by Genemed Synthesis, Inc. (San Francisco, CA). These S2 fragment peptides were synthesized as 25-mers with 11-amino acid overlap with the following amino acid sequences (20): S2-1 SAEREKEMASMKEEFTRLKEALEKS,S2-2 FTRLKEALEKSEARRKELEEKMVSL, S2-3 RKELEEKMVSLLQEK NDLQLQVQAE, S2-4 KNDLQLQVQAEQDNLADAEERCDQL, S2-5 LADAEER CDQLIKNKIQLEAKVKEM, S2-6 KIQLEAKVKEMNERLEDEEEMNAEL, S2-7 LEDEEEMN AELTAKKRKLEDECSEL, 2-8 KRKLEDECSELKRDIDDLELTLAKV, S2-9 IDDL ELTLAKVE KEKHATENKVKNL, S2-10 KHATENKVKNLTEEMAG LDEIIAKL, S2-11 MAGLDE IIAKLTKEKKALQEAHQQA, S2-12 KKALQ EAHQQ ALDDLQAEEDKVNTL, S2-13 LQAEEDKVNTLTKAKVKLEQ QVDDL, S2-14 KVK LEQQ VDDLEGSLEQEKKVRMDL, S2-15 LEQEKKVRMDLER AKRKLEGDL KLT, S2-16 KRKLEGDLKLTQESIMDLENDKQQL, S2-17 IMDLENDKQ QLDERLKKKDFELNAL, S2-18 LKKKDFELNALNARIEDEQALGSQL, S2-19 IEDEQALGSQLQ KKLKELQARIEEL, S2-20 LKELQARIEELEEELESERTARAKV, S2-21 LESERTAR AKVEKLRSDLSR ELEEI, S2-22 RSDLSRELEEISERLEEAGGATSVQ, S2-23 LEEA GGATSVQIE MNKKREAEFQKM, S2-24 NKKREAEFQKMRR DLEEATLQ HEAT, S2-25 LEEATLQHEATAAALRKKHADSVAE, S2-26 LRKKHADSVAEL GEQIDNL QRVKQK, S2-27 QIDNLQRVKQKLEKEKSEFKLELDD, S2-28 EKSEFKLELD DVTSNMEQIIKAKAN, S2-29 NMEQIIKAKANLEKMCRTLEDQMNE, S2-30 MCRTLED QMN EHRSKAEETQRSVND, S2-31 KAEETQRSVNDLTSQRAKLQTENGE, S2-32 ETQR SVNDLTSQRAKLQTENGELSR.

### Protein Kinase A (PKA) Assay

H9c2 rat cardiac myoblast cell line (ATCC) (1 x 10^7^) were plated in T75 cell culture flasks overnight at 37^0^ at 5% CO_2_ for use in the protein kinase A(PKA) assay. H9c2 has been demonstrated to express the beta adrenergic receptor(51). Serum samples were incubated with the H9c2 cells at a 1:100 dilution in a final volume of 15 mL of serum-free medium for 1 hour before reactions were stopped by an ice-cold PBS (10 mL) rinse. The cells were mechanically dislodged from the flasks, centrifuged, and solubilized in 0.3 mL of protein extraction buffer before homogenization. Each pellet was homogenized with chilled homogenizer for 30 seconds, on ice. PKA activation in H9c2 cells was measured using SignaTECT cAMP-dependent protein kinase assay system (Promega) according to the manufacturer’s instruction. The specific activity of the enzyme in picomoles per minute per microgram for each sample was calculated and results were presented as percentage above the basal PKA rate (20,52).

### Human Hybridoma Production

Human hybridoma production was performed by routine methods in our laboratory for creating and maintaining hybridomas and producing human monoclonal antibodies (mAbs) (45,53–55). Peripheral blood was obtained from a 54-year-old female diagnosed with DCM. Peripheral blood mononuclear cells were separated from whole blood by Histopaque-1077 (Sigma) and stimulated for 1 week with pokeweed mitogen (2 μg/mL) in Iscove’s Modified Dulbecco’s Medium (IMDM) containing 10% human AB serum, 1µL/mL gentamycin, and penicillin streptomycin (1mL/50mLs). Cells were washed three times in Iscove’s Modified Dulbecco’s Medium (IMDM) (serum-free) and fused with HMMA2.11TG/0 cells (human/mouse myeloma cell line). The HMMA cell lines have been described by their inventor Dr M R Posner(56). HMMA 2.11TG/O was the fusion partner with the human lymphocytes purified by ficoll-hypaque gradient from the peripheral blood of the myocarditis patient. HMMA cells were added to the lymphocyte suspension in a 1:1 ratio and centrifuged together at 1000rpm for 7 minutes. Three milliliters of 35% polyethylene glycol (PEG-1000) in IMDM pH 7.8 were dripped down the side of the tube. The cell mixture was gently agitated and centrifuged at 900rpm for 2 minutes. The cells were then “rested” for 6 minutes at 30^0^C. Forty milliliters of IMDM containing 10% FBS was gently dripped down the side of the tube. The tube was centrifuged at 900 rpm for 8 minutes and the cell pellet was re-suspended in IMDM containing 10% FBS, gentamycin, and penicillin streptomycin. The cells were plated in 24-well tissue culture plates at 1×10^6^ cells/mL and 0.5mL/well was placed into each well and incubated overnight. The next day, 0.5mL HAT (hypoxanthine, aminopterin, and thymidine) medium was added to each well. On day 3 after the fusion, 0.5mL fresh HAT medium was added to each well. On days 5 and 10, 1mL fresh HAT medium was added to each well. Starting on day 12, the medium was switched to HT (hypoxanthine and thymidine) and 1mL fresh media was added every 3-5 days as hybridoma growth was monitored. Culture fluid from wells positive for hybridoma growth was screened in the enzyme-linked immunosorbent assay (ELISA) against antigens CM and β1AR. Cloning of hybridomas was achieved by limiting dilution and rescreening with antigens and was performed two times. Established clones were maintained in IMDM containing 20% fetal bovine serum (FBS) eventually reduced to 10% FBS for final fluid collection for study. Assays were compared to the media control containing FBS. Positive clones were identified for immunoglobulin isotype and concentration using standard curves and isotype-specific secondary antibodies.

### Monoclonal Antibody 2C.4

Human hybridoma cell lines were selected by supernatant reactivity to βAR and PKA activity by ELISA, cloned twice, stored, and maintained as described above. The human mAb used in this study included 2C.4 which was stored as culture fluid at 4^0^C. Clone 2C.4 was screened against human cardiac myosin and the β-adrenergic receptor (βAR) enriched membranes (Perkin Elmer) by ELISA, and by PKA activity. Our process resulted in several clones with strong reactivity to βAR1 and βAR2 (**Supplementary figure 6**). We selected clone 2C.4 because it was highly cross-reactive with human cardiac myosin, had the highest reactivity to both βAR 1 and 2, and activated PKA in the ATCC H9c2 heart cell line.

### Gene Expression in Treated and Untreated H9c2 (ATCC) Heart Cell Line

The rat cardiac myoblast/primary heart cell line H9c2 (ATCC) was described by ATCC as H9c2(2–1) CRL-1446 as: “a subclone of the original clonal cell line derived from embryonic BD1X rat heart tissue that exhibits many of the properties of skeletal muscle. This cell line is recommended for cardiovascular disease research.” The H9c2 cell line was treated in triplicate for 30 minutes with 1:100 dilution of sera from myocarditis/DCM patients (n=2) or healthy subjects (n=2), left untreated, or treated with 0.882 ug/mL of mAb 2C.4. Following treatment, RNA was isolated from individual wells using TRIzol. RNA concentrations were quantified using a Qubit fluorometer, and mRNA libraries were generated using 500ng of each RNA sample and Illumina’s TruSeq Stranded mRNA Library Prep Kit. Bulk PBMC RNA-seq was performed on the Illumina HiSeq 3000 (Illumina Inc., San Diego, CA) in the OMRF’s Clinical Genomics Center following Center procedures. Samples were sequenced using 75bp paired-end reads with 10 samples per lane, which yielded ∼30 million reads per sample. Post-sequence reads were quality filtered and trimmed for Illumina adapters using Trimmomatic v0.35 (57). Resulting reads were pseudo-aligned to coding regions of the Rnor_6.0 genome (release 101) genome using Kallisto v0.44.0 (58) with the following options: bias enabled, 50 bootstraps. Expression values for transcripts were measured in transcripts per million reads sequenced (TPM). Gene annotation was performed via the R package biomaRt (59). To perform differential expression (DE) analyses, expression values were summarized at the gene level to transcript-length adjusted, library-size scaled counts per million (CPM) with the R package tximport (60). Differential expression was calculated using the empirical Bayes approach implemented in the R package DESeq2 (61). Significantly DE genes had an absolute value log2 fold change ≥1.

### Pathway Analyses

To characterize pathways, we utilized a commercial software package, Ingenuity Pathway Analysis (IPA) to intersect DE genes with known biological functions or pathways, maintained in the IPA database, a collection of nearly 5 million experimental findings manually curated from either literature or third-party databases. Detailed information can be found at http://qiagen.force.com/KnowledgeBase/. Enriched pathways are identified via the Canonical Pathways function. The significance of the overlapping genes from DE genes and genes within the enriched pathway is reported as a p-value, calculated by the right-tailed Fisher’s Exact Test. In addition to a p-value, Canonical Pathways reports the percent overlap (given as a ratio) and a z-score, a measure of activation or inhibition of a pathway based on the observed and predicted magnitude and direction of fold changes in overlapping genes. Calculations of z scores have been described in detail (62). Here, a pathway was defined as enriched if its IPA Z score was above 1 or below -1 and had a significant p-value (<0.05).

### Confirmation of RNAseq by PCR

Eight genes were selected from our RNA sequencing result for confirmation by PCR via an independent experiment with parallel design to our RNA sequencing experiment. As described above, H9c2 rat heart cells (ATCC) were incubated with patient (n=6) and healthy subject (n=4) sera (1:100) for one hour in medium and washed with ice-cold PBS. Cells were immediately lysed with 1ml TRIzol (Invitrogen) and stored at -80°C. RNA was isolated by chloroform extraction according to manufacturer’s (Invitrogen) instructions. RNA cleanup was performed by passing isolated RNA through silica columns (RNA Clean & Concentrator kit, Zymo Research.). cDNA was reverse transcribed from 0.35ug RNA with OmniScript Reverse Transcriptase (Qiagen), diluted 1:10, and used directly in 25ul SYBR Green (QuantiTect SYBR Green PCR Kits, Qiagen) PCR reactions at a final dilution of 1:125. Pre-designed and validated primers (KiCqStart SYBR Green Primers, Millipore Sigma) were used at a final concentration of 0.5 µM. 40 cycles of PCR were carried out on ABI 7500 RT-PCR system, and CT threshold set in the linear range of amplification, and a single amplicon was verified by a single product present in the dissociation curve. Non-specific amplification was controlled through control reactions lacking template, and with no-RT controls. Log_2_ fold changes were calculated using the 2^-ΔΔCt^ method (63), and statistical significance between groups was determined using a two-tailed unpaired t-test.

### Statistical Analyses

For variables including anti-CM titer and PKA activation, when the normality of the distribution was rejected, a nonparametric Kruskal-Wallis test was used for comparisons of medians among 3 or more groups, followed by post-hoc testing using unpaired Mann-Whitney *U* tests with a Bonferroni-adjusted alpha level. One-way ANOVA with a Tukey’s post-hoc multiple comparison test was used to compare more than 3 means when normality was satisfied. Correlation between CM titer and echo LVEF was determined by using the Spearman’s rank correlation test. A 2-sided p-value less than or equal to 0.05 was considered statistically significant, unless otherwise specified, where a more stringent, or conservative, alpha level less than 0.05 was used to define significance due to multiple comparisons. Bar graphs reflect the mean ± SEM, while the dot plots include a horizontal line drawn at the median value for each subgroup.

### Reporting Sex and Gender Based Analyses

Although we evaluated both sexes in our study, more males (n=27) than females (n=14) are found to have myocarditis and the number of the non-recovered patients was small (n=6) with 3 males and 3 females in the non-recovered cases studied. Our previous study of this cohort did compare males and females and is published (6). Males were dominant for biomarkers of non-recovery such as Il-17A and IL-6, and in another study, elevated ST-2 dominated in males with myocarditis. Here the comparison of cardiac myosin Ab titers in males vs females was 25 males to 12 females with no statistical differences observed.

### Study Approval

The use of subject blood and data required for our studies was approved by Institutional Review Boards at OUHSC and Mayo Clinic. Written informed consent was obtained from participants before study initiation according to the Declaration of Helsinki with regard to scientific use. Peripheral blood was obtained from healthy subject subjects who were laboratory volunteers at OUHSC or were healthy donors from the Oklahoma Blood Institute in Oklahoma City, Oklahoma, USA.

### Data Availability

Sequencing data can be accessed through NIH National Center for Biotechnology Information under BioProject accession PRJNA1025010.

### Author Contributions

JMM and CES contributed equally to this work. MWC, JMM, and LTC conceived and designed the study. JMM, MWC, and GW developed the methods and JMM, MWC, CES, and LG analyzed and interpreted the data. JMM, CES, MWC, KA, LG, GW, CM, PG, DF, LTC, and SS wrote, reviewed, and revised the manuscript.

## Results

### Nonrecovered myocarditis patients at baseline show significantly higher cardiac myosin (CM) and CM peptide autoantibodies than those who recover

We hypothesized that CM AAbs may be a prognostic biomarker for nonrecovery of LVEF in myocarditis and dilated cardiomyopathy (DCM) and tested this hypothesis in a longitudinal study of myocarditis/DCM patients where recovery was defined as reaching ≥50% LVEF by one year after onset. Both recovered and nonrecovered myocarditis/DCM patients were analyzed for IgG reactivity to human CM, and to 32 overlapping synthetic peptides of the S2 hinge-region fragment of CM (CM peptides). Sera was collected at initial clinical presentation at baseline, no more than 6 months past onset, and tested by ELISA for IgG reactivity to CM and CM peptides. In myocarditis patients, elevated titers of serum CM AAbs trended towards significance (p=0.0533) in baseline samples compared to healthy subjects (**Figure 1A**). However, highly significant elevated CM AAb titers were found in our cohort of nonrecovered patients (n=6) (**Figure 1B**, p=0.0096) compared to normal healthy subjects. Elevated CM AAbs in the nonrecovered group suggest that CM AAbs may be a biomarker of disease progression similar to AAbs in other autoimmune diseases (64) and may be useful for prognosis of outcomes in myocarditis. We also analyzed AAb titers to peptides from the S2 hinge region of CM, and we noticed that in the nonrecovered group, IgG reactivity appeared significantly higher than in the recovered group. We hypothesized that reactivity to all peptides may be more important than any single one. To test this, we averaged all patient IgG reactivity for each given peptide and compared the means of all peptides measured in the recovered group with the nonrecovered group. We found that baseline sera IgG reactivity to S2 peptides was significantly elevated in nonrecovered myocarditis/DCM patients when compared with recovered patients (**Figure 2C**, p<0.0001), suggesting that measuring IgG reactivity to CM peptides may be important in determining disease outcomes. To study the relationship of CM AAb titers to outcomes, we correlated LVEF with titers in our patient cohort. LVEF did not significantly correlate with CM AAb titers in patients who recovered LVEF (**Supplementary Figure 1**, r=-06274, p=0.7559), however, the subset of myocarditis patients (n=6) who did not recover LVEF (nonrecovered) were found to have high CM AAb titers which correlated with low LVEF (**Figure 2D**, r= -0.8827, p=0.0167). Removal of one apparent outlier (patient with AAb titer 6400) resulted in an even stronger correlation (**Supplementary Figure 2**, r = -0.9487, p= 0.0333). Thus, CM AAbs negatively correlated with LVEF in patients who did not recover, suggesting a functional role of CM AAbs in progression of myocarditis.

**Figure 1.**
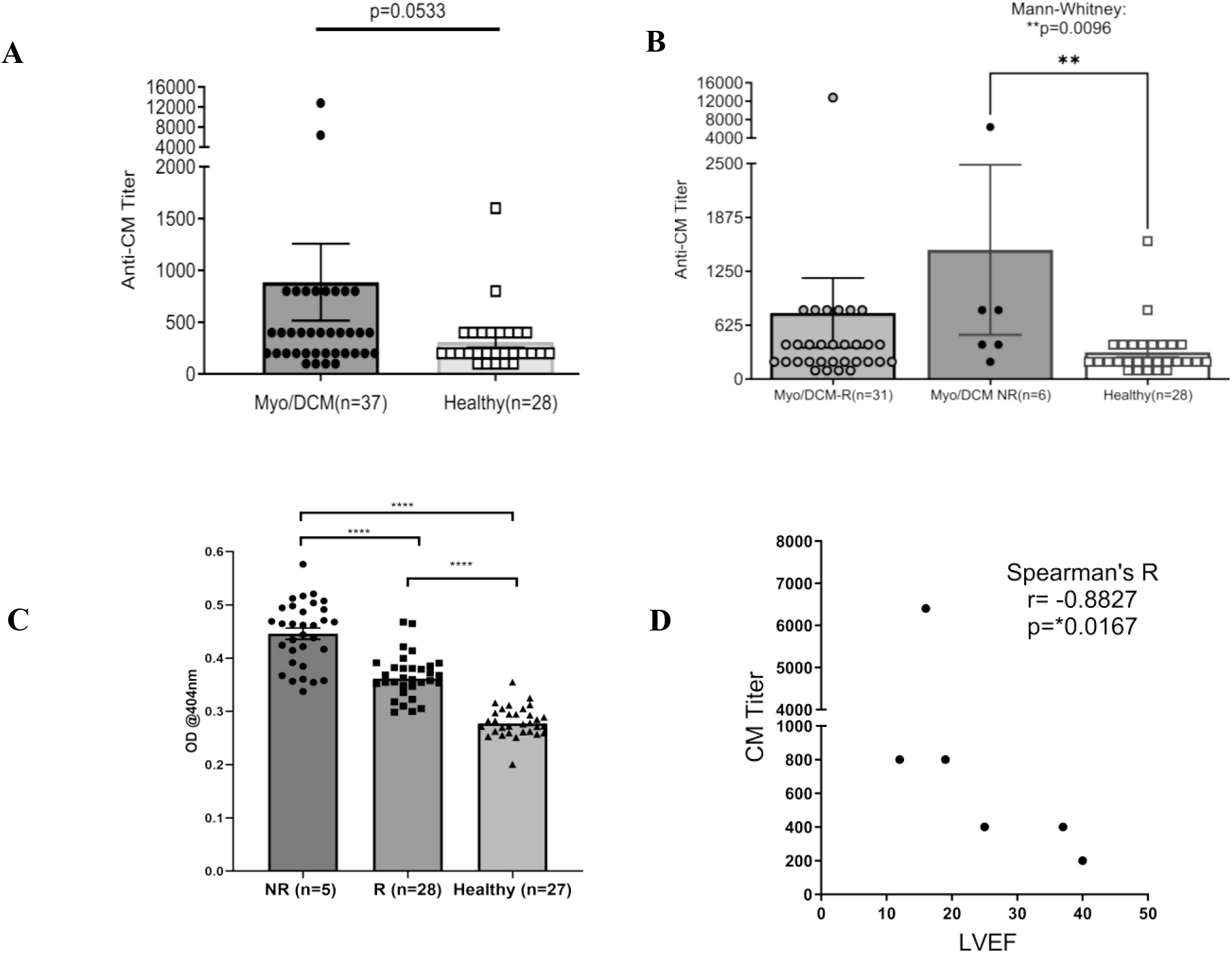
Baseline autoantibodies to CM peptides are significantly elevated in nonrecovered (NR) myocarditis patients compared with recovered (R) patients. Sera taken at initial clinical presentation from recovered and nonrecovered myocarditis were reacted with whole CM molecule or 32 overlapping peptides of CM in ELISA. CM titers trended towards significance between all patients and healthy subjects **(A)**. Nonrecovered patients had significantly elevated titers to CM compared to healthy controls **(B** In **(C)**, Nonrecovered (n=6) myocarditis patients have significantly elevated autoantibodies to CM peptides when compared with recovered (n=28) patients and healthy subjects (n=27). In **(C)**, each datapoint is the average serum IgG immunoreactivity (OD @ 1:100 serum dilution) with all 32 S2 CM fragment peptides for each individual (i.e. each black dot is the average from all patients for one peptide). Serum IgG titers to CM correlated with lower ejection fraction in nonrecovered patients, but not in recovered patients **(D** and **Supplement Figure 1**) NR = Non-recovered, R = recovered

**Figure 2.**
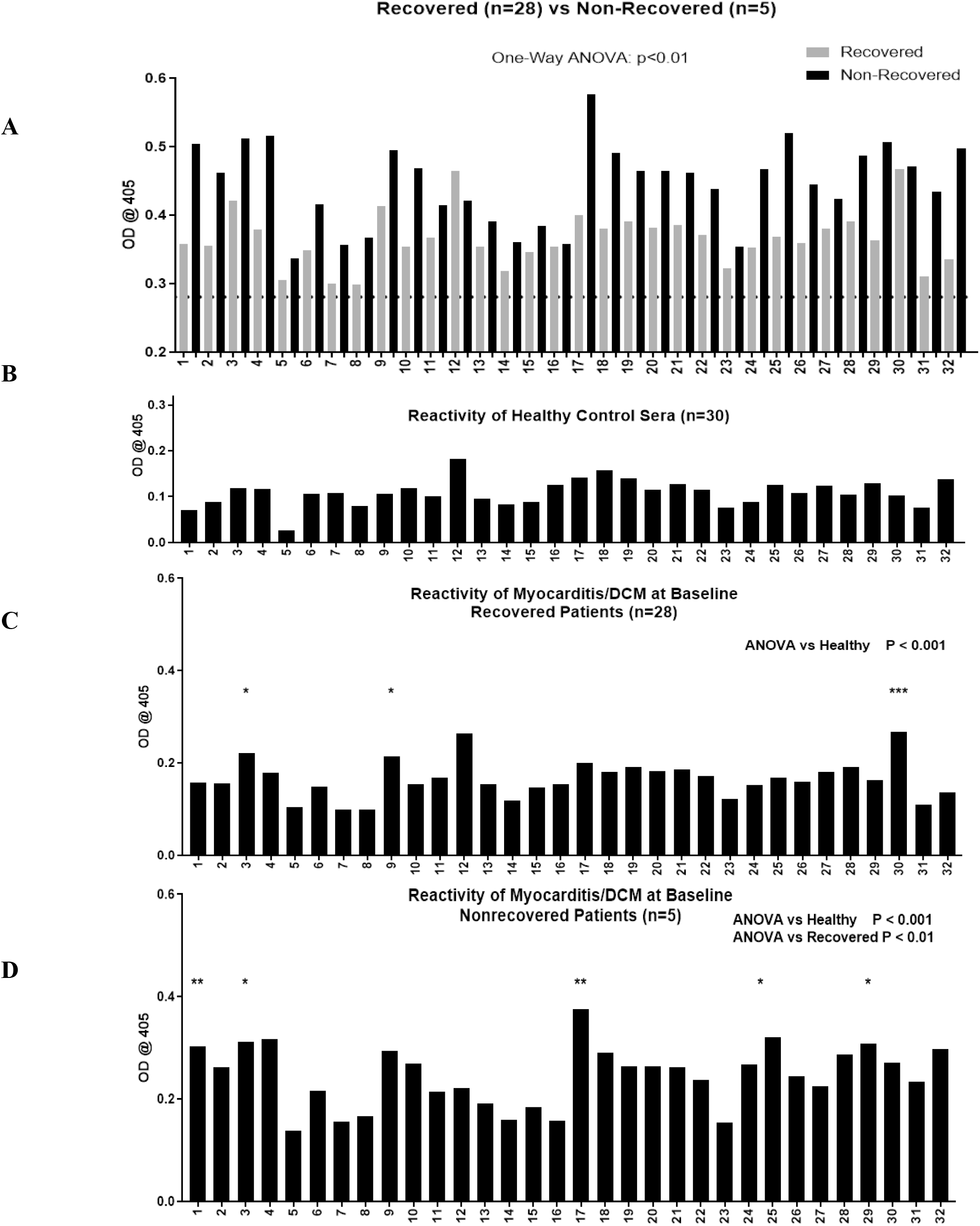
Autoantibodies (AAbs) to CM peptides are elevated in nonrecovered patients, and AAbs recognize unique CM peptide epitopes in nonrecovered patients. Overlapping peptides from the S2 region of CM were tested by ELISA for reactivity with serum IgG in healthy (n=30, **Panel B**), recovered (n=28, **Panel C**), and nonrecovered patients (n=5, **Panel D**). Recovered and nonrecovered are shown in the top Panel (**A**), dotted black line indicates mean healthy subject (n=30) serum IgG antibody reactivity at mean OD of 0.2809 ± 0.03. OD values normalized to healthy are plotted in **B**, **C** and **D**. One-way ANOVA showed groupwise differences between healthy compared with recovered (panel C, p <0.0001) and with nonrecovered (panel D, p <0.001). Post-hoc analysis showed that in recovered patients compared with healthy control serum IgG, peptides S2-3 (p <0.01), S2-9 (p < 0.01), and S2-30 (p <0.0001) reached significance after multiple test correction. <u>IgG AAbs to CM peptides</u> <u>specific to nonrecovered patient sera were S2-1 (p < 0.001), S2-17 (p < 0.001), S2-25 and S2-</u> <u>29</u> (both p <0.01) reached significance above healthy subjects. ANOVA showed groupwise differences between recovered and nonrecovered patients (panel D, p < 0.01).

We sought to identify unique CM epitopes of nonrecovery in myocarditis patients. OD measurements for all individual S2 hinge region peptides are shown in **Figure 2A**, where grey bars show the average OD for all recovered patients, and black bars correspond to nonrecovered patients. The S2 hinge region was chosen in our studies because it was demonstrated to be highly antigenic in previous studies (20), likely due to its proteolytic sensitivity (65). The dashed line indicates mean healthy subject IgG reactivity (optical density at 0.2809 ± 0.03) (**Figure 2A**). Groupwise analysis of all 32 peptides showed significant differences between healthy compared with both recovered and nonrecovered patient groups (One-way ANOVA, both p < 0.0001), and post-hoc analysis revealed several individual peptides that reached significance after multiple test correction (**Figure 2C and D)**. Further, recovered patients compared with nonrecovered was also significant (One-way ANOVA p<0.01). Therefore, significantly elevated IgG specific for S2 CM peptides at the time of clinical presentation suggest that they may be an important prognostic biomarker for nonrecovery in human myocarditis/DCM and warrant further investigation in a larger cohort.

### Myocarditis/DCM sera IgG activates protein kinase A (PKA) significantly higher than healthy sera

We previously demonstrated that autoantibodies specific for cardiac myosin cross-react with βARs and activate protein kinase A (PKA) (14,20). Activation of cyclic AMP-dependent PKA is a key step in cardiomyocyte contraction, and overstimulation of this pathway may lead to cardiomyocyte apoptosis and contribute to dilated cardiomyopathy (14). CM AAbs bind directly to both βAR1 and βAR2, and signaling through PKA can be inhibited pharmacologically by βAR antagonists and by pre-incubation of AAbs with soluble βAR1, βAR2 and human cardiac myosin (14,20). In **Figure 3** we demonstrate that myocarditis/DCM sera activate PKA signaling in a rat heart cell line (ATCC H9c2) using a P^32^ detection assay. Myocarditis/DCM sera at each time point (baseline, 1, 6, and 12 months) activated PKA significantly higher than healthy sera (**Figure 3**), however no significant differences in PKA activation between recovered and nonrecovered patients were observed (**Figure 3**). PKA activation was significantly (p=0.0061) higher at the myocarditis baseline blood draw compared to the 12-month blood draw, perhaps related to recovery in most patients by 12 months. In a small myocarditis patient sample set we verified that, as in our previously published work (14,20), PKA signaling in our current cohort was indeed due to IgG mediated signaling (n= 3, **Supplementary Figure 3**). Incubation of myocarditis sera with propranolol diminished PKA activation, and anti-IgG-coated beads diminished PKA activity in a similar magnitude while BSA-coated beads demonstrated no effect (**Supplementary Figure 3**). In our previous studies, PKA signaling was blocked >80% by co-incubation of CM AAbs with antigens CM, β1AR, and β2AR, demonstrating cross-reactivity of the CM AAbs with βAR (20). In context with previously known properties of CM AAbs, our present results suggest PKA signaling induced by myocarditis/DCM sera reflects βAR activation by AAbs specific for both CM and the βAR.

**Figure 3.**
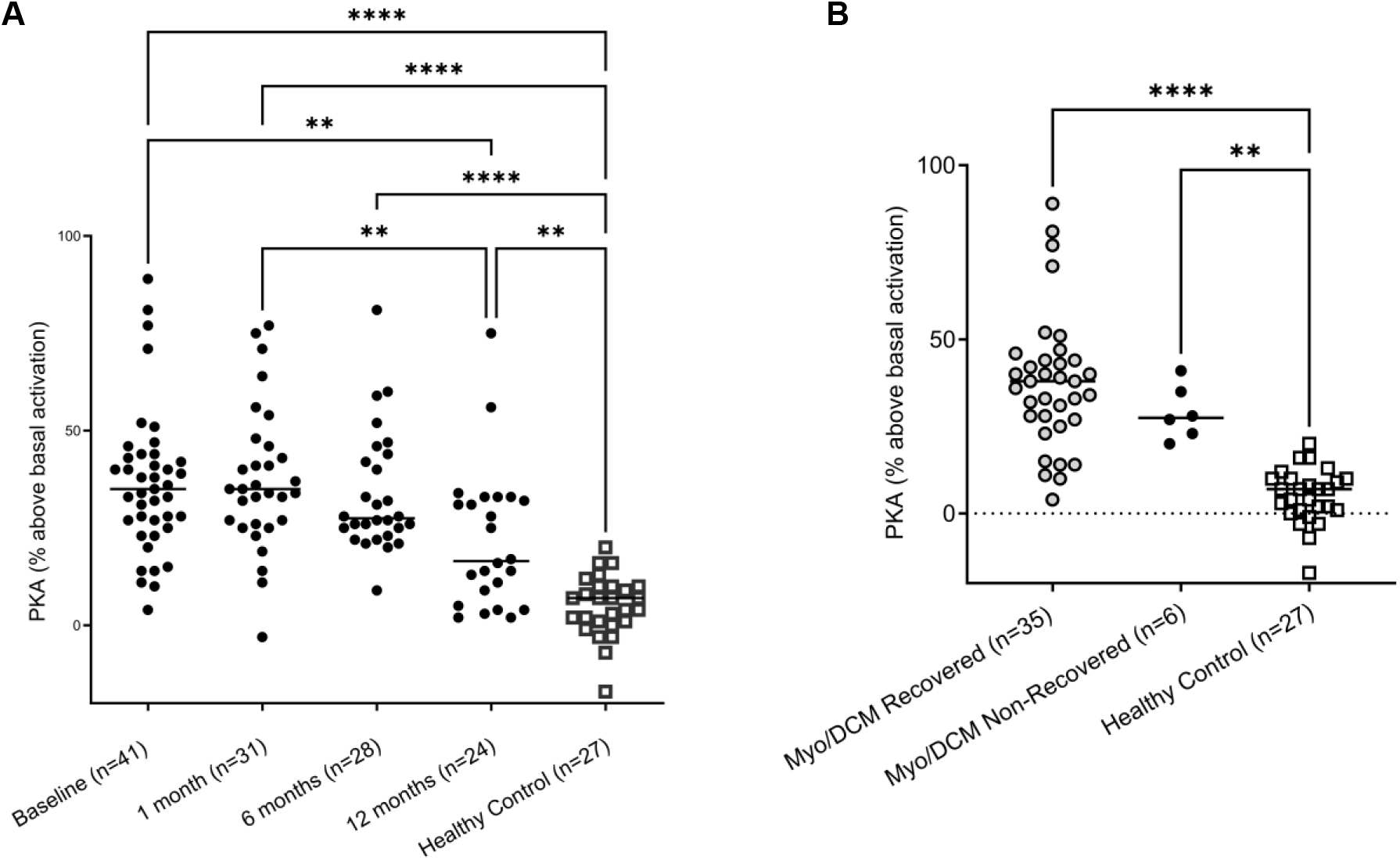
Myocarditis/DCM sera signal PKA. H9c2 (ATCC) was treated with baseline myocarditis/DCM sera (1:100 dilution) Levels of PKA signaling in H9c2 cells are compared to healthy subject levels. PKA signaling is significantly elevated in both recovery and non-recovery at the baseline time point. Recovered is shown in One-way ANOVA compared with healthy sera with Tukey’s multiple comparison test ****p<0.0001, Nonrecovered vs Healthy sera: **p=0.0023. Although myocarditis sera strongly signal PKA, there is no difference in the recovered vs nonrecovered.

### Human myocarditis sera promote a profibrotic and proinflammatory transcriptome in H9c2 cells

To investigate other potentially harmful effects of CM AAbs, we investigated the transcriptome of untreated H9c2 cells compared with H9c2 cells treated with human myocarditis vs healthy sera. Treatment of H9c2 cells with myocarditis-derived human mAb 2C.4, myocarditis-DCM patient sera, and sera from healthy subjects induced differential expression of 1296, 472, and 470 genes, respectively, relative to untreated H9c2 cells (basal). Over half (259/472, 55%) of differentially expressed (DE) genes in H9c2 cells treated with myocarditis-DCM patient sera were also DE in mAb-treated cells (**Figure 4B**), and nearly all (238/259) of these were concordant in sign (i.e. positive in both analyses, or negative in both analyses). **See Figure 4B**.

**Figure 4:**
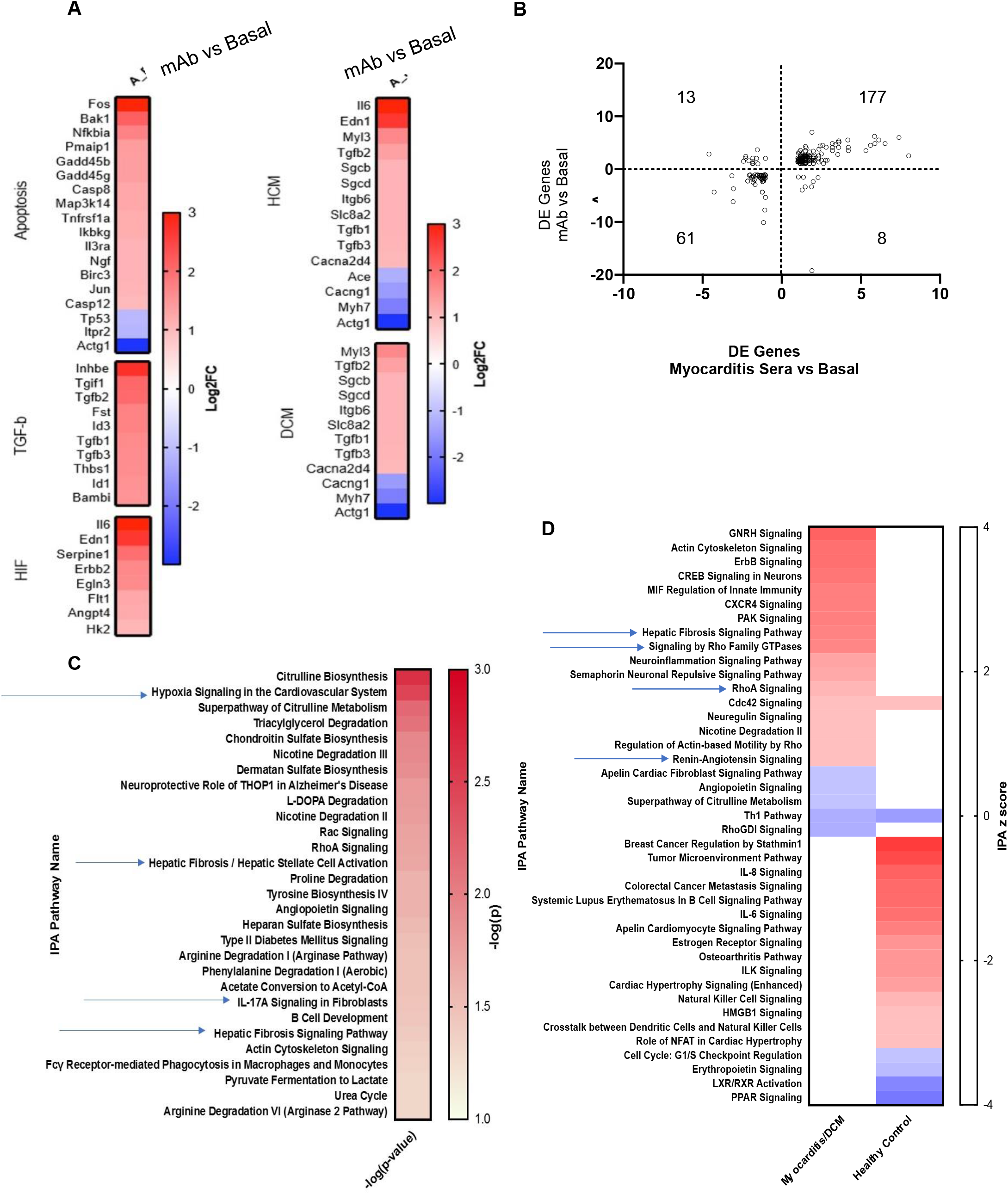
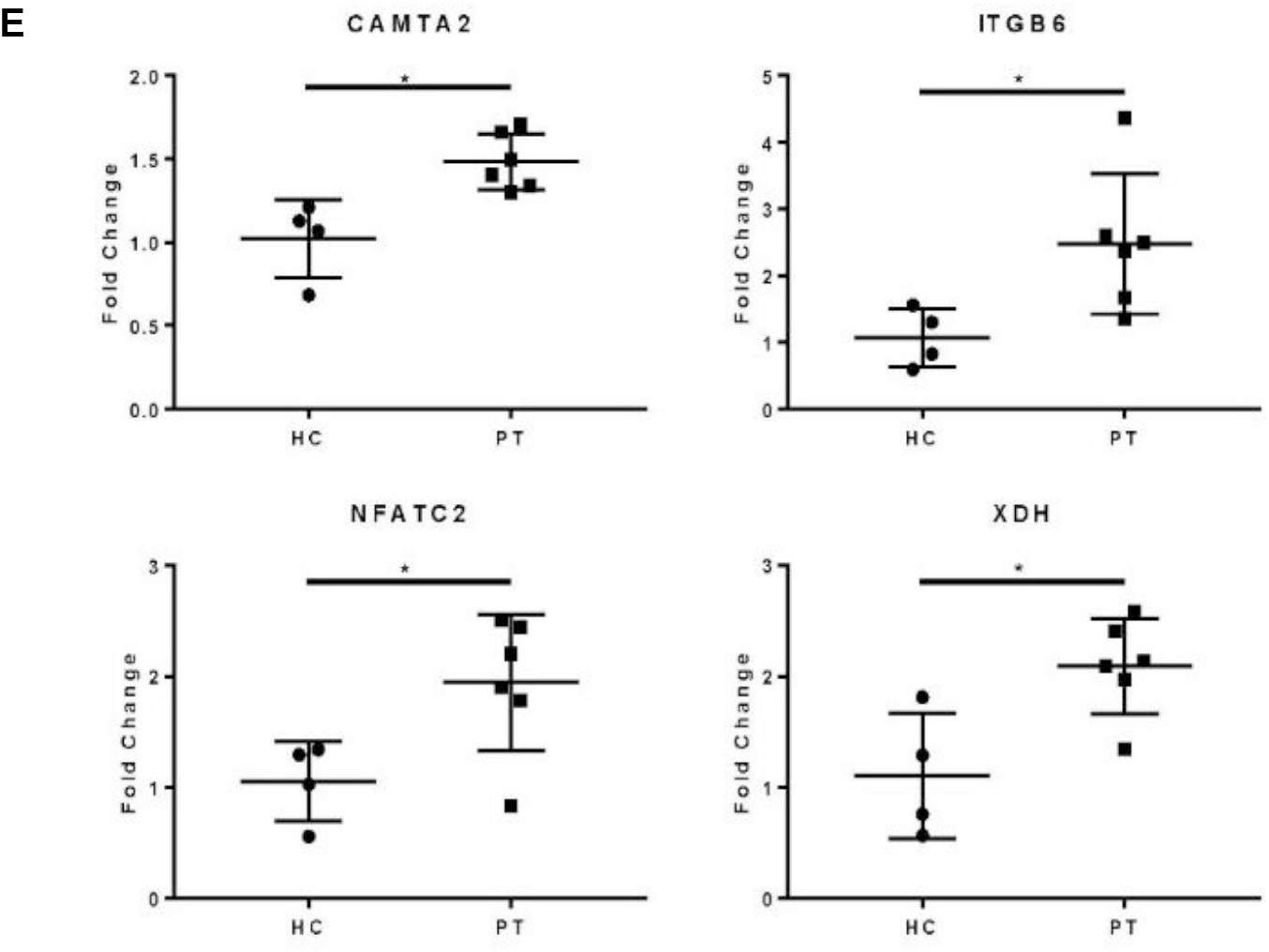
Myocarditis-derived human mAb 2C.4 and myocarditis-derived sera promote fibrotic transcriptome in H9c2 cells. (**A**) KEGG Analysis of DE genes revealed that myocarditis derived human mAb or sera compared to basal treated H9c2 heart cells induced expression of genes associated with TGFβ expression/fibrosis, apoptosis, HCM or hypertrophic cardiomyopathy and DCM or dilated cardiomyopathy. (**B**) DE genes in mAb/sera vs. basal treatments. Log2FC values of DE genes showed high concordance (same direction; ie positive or negative direction) in mAb/sera treatments: DE genes in upper-right (177) vs lower left (61) quadrants. (**C**) Ingenuity Pathway Analysis (IPA) of DE genes identified in myocarditis(n=2) vs healthy (n=2) serum treated H9c2 cells shows enriched fibrosis/hypoxia/apoptosis pathways. Red positive z score; blue negative z score indicates DE direction. Arrows indicate fibrosis related pathways identified. (**D)** IPA of DE fibrosis related genes in myocarditis (upper left) but not in healthy (*lower right*) treatments p < 0.05. Arrow**s** indicate fibrosis related pathways identified. (**E**) Myocarditis(n=6) vs healthy sera (n=4) treated H9c2 cells were examined by PCR: Fibrosis/Heart Failure genes were upregulated in 6 myocarditis vs 4 healthy patient(s) (PT) (CAMTA2, ITGB6, NFATC2, and XDH) Data analyzed with ΔΔCT method; β-actin reference gene. A gene was considered DE if it demonstrated Log_2_-fold change > or < 1 (red and blue, respectively).

DE genes were used for pathway analysis in both Ingenuity Pathway Analysis (IPA) and KEGG (Kyoto Encyclopedia of Genes and Genomes) databases to investigate altered pathways in DE genes in H9c2 cells treated with myocarditis-derived human mAb and sera. H9c2 cells treated with the myocarditis-derived human mAb 2C.4 show enrichment of genes associated with hypoxia (HIF), fibrosis (TGF-beta 1) and apoptosis, as well as genes associated with hypertrophic and dilated cardiomyopathies (**Figure 4A**). Pathways enriched (IPA; z score >1 or <-1, p<0.05) from treatment of H9c2 cells with myocarditis-DCM patient sera relative to basal were strikingly different from those enriched from treatment with sera from healthy subjects (**Figure 4D**). Hepatic fibrosis signaling was enriched only in H9c2 cells treated with myocarditis sera. Specific genes DE in myocarditis sera-treated H9c2 cells were not DE in H9c2 cells treated with healthy subject sera (256/472) including fibrosis-associated genes CACNA1S, CCN2, COL5A3, FTL, MYLK3, NFKBIE, and TGFBR3. Canonical pathway analysis (IPA) additionally revealed two pathways related to the pathogenesis of myocarditis, including ‘Hypoxia Signaling in the Cardiovascular System’ and ‘IL-17A Signaling in Fibroblasts’ with unadjusted p-values <0.05; however, these two enriched pathways did not reach significance due to low value of z scores and likely due to the very small sample size used in these hypothesis-generating pilot experiments (**Figure 4C**). We compared enriched pathways (IPA) between myocarditis sera-treated and healthy subject sera-treated H9c2 cells and found fibrosis pathways significantly (p<0.05) elevated in the myocarditis sera treated H9c2 cells but not in the healthy subject sera treated H9c2 cells (**Figure 4D**).

An independent RT-PCR experiment was performed with parallel design to our RNAseq experiment which compared H9c2 cells treated with serum from 6 different myocarditis patients with 4 healthy subjects, and PCR data was analyzed using the ΔΔCT method (63) with β-actin as the reference control gene. By PCR, an additional 4 genes involved in heart failure pathogenesis were found with their gene expression significantly elevated in H9c2 cells after treatment with the myocarditis patient sera but not with sera from healthy subjects. CAMTA2, ITGB6, NFATC2, and XDH were all significantly elevated and differentially expressed after treatment with the myocarditis sera (p= <0.05) (**Figure 4E**). Several other genes tested by PCR (NLRP3, PTPRQ, CFH, and SDK2) were not elevated over healthy subject gene expression values (not shown). ITBG6 (Integrin Beta 6) was elevated in both the RNA seq analysis as well as the PCR expression studies. Overall, these data suggest that AAbs against CM, including a myocarditis-derived human mAb and myocarditis sera can promote gene expression of fibrosis, apoptosis, and hypoxia genes and pathways in H9c2 cells.

### Human myocarditis derived mAb 2C.4 induces fibrosis pathways in H9c2 cells

We hypothesized that CM AAbs would have other important effects on H9c2 cells in addition to PKA activation. To test this hypothesis, we performed gene set enrichment analysis (GSEA) (66,67) on mAb 2C.4-treated H9c2 cells compared with isoproterenol (ISO, synthetic βAR agonist) treated H9c2 cells. The visualization suite R package enrichplot (68) was utilized to aid in interpretation of the results (**Figure 5**). From the entire list of enriched pathways, we used the function emapplot() from the R package enrichplot to visualize all significantly enriched pathways with a positive enrichment score, and to cluster pathways into functional units (69) (**Figure 5**, top panel). We then identified two pathway clusters, one related to known interactions of β-adrenergic receptors, and the second related to fibrosis genes differentially expressed in H9c2 cells. (**Figure 5, top panel, clusters A, B**). The genes within the leading edge were visualized using the function cnetplot() from the R package enrichplot (68) (**Figure 5, panels A & B**) to determine whether pathways are independently enriched, or merely represent child-parent relationships. **Figure 5** panels **A** and **B** show that most pathways are enriched from unique sets of genes, with a few redundant (child-parent) pathways.

**Figure 5:**
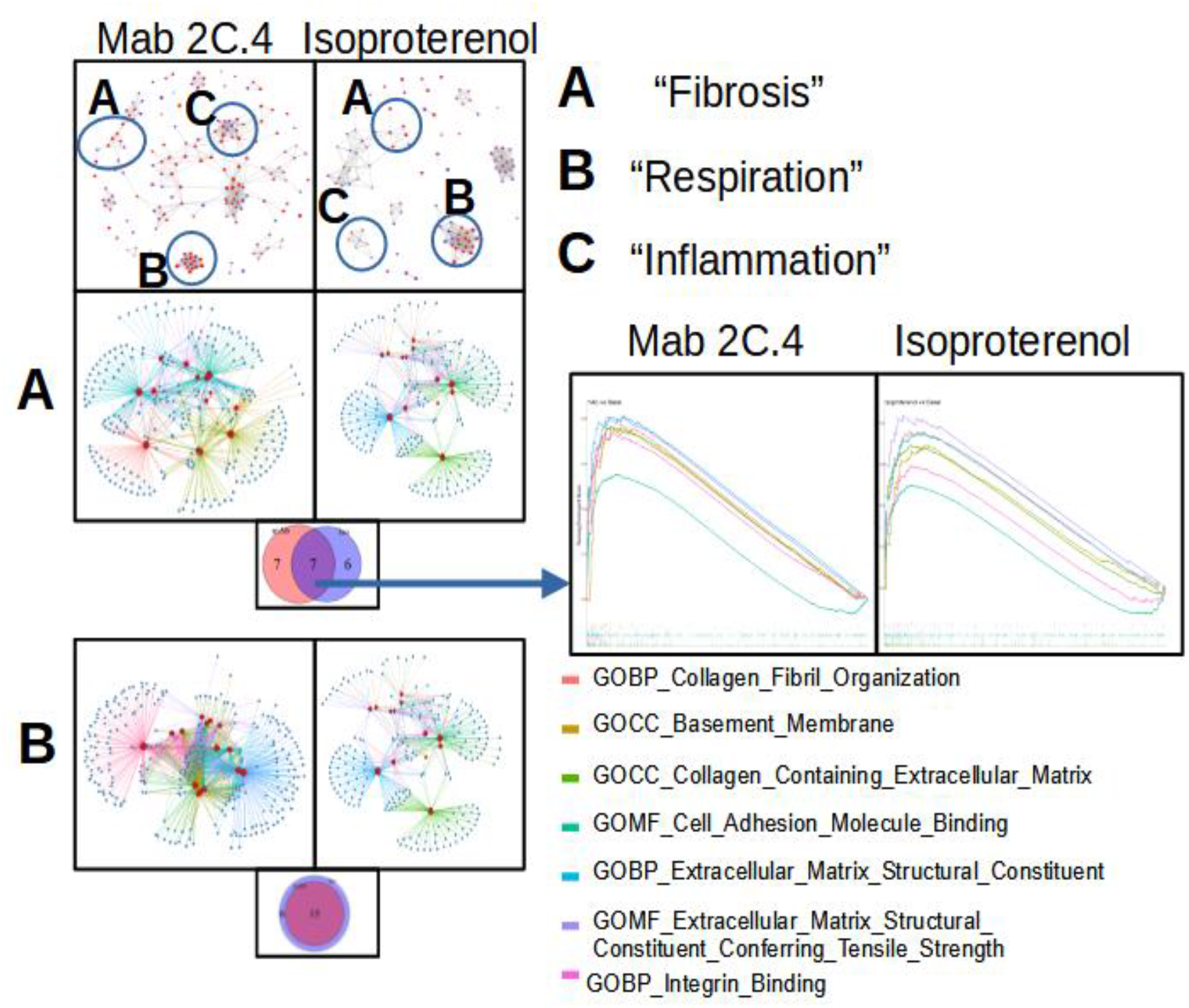
Myocarditis-derived mAb 2C.4 induces fibrosis-related pathways in heart cells. Heart (H9c2) cells treated with human mAb 2C.4 or β-adrenergic agonist isoproterenol (ISO) underwent RNA seq subsequent to pathway analysis by GSEA. Functional units (FU) visual-ized with emapplot() from enrichplot (top panel) were related to fibrosis, respiration, and in-flammation identified from upregulated, significantly enriched pathways (top panel). (**A**) Fi-brosis pathways were identified from mAb vs ISO treatments of cells. Genes from leading edge plotted with enrichplot() ruled out redundant pathways. Unique/shared pathways between ISO and 2C.4 are shown in Venn-diagram (inset) and elevated fibrosis pathways compared (arrow). GSEA plots for seven shared fibrosis pathways reveal similar mAb 2C.4 and ISO pathways identified in H9c2 cells suggesting shared fibrosis signatures. (**B)** Myocarditis-derived mAb2C.4 induced many respiration pathways similar to ISO. (**C**) ISO enrichment maps re-vealed cluster of T-cell activation inflammatory pathways (top right panel) which had no similarity to ISO, while strong similarities/overlap between ISO and mAb 2C.4 were evident in respiration and fibrosis pathways.

It is well established that β-adrenergic stimulation causes an increase in respiration (70–74) and mAb 2C.4 was selected on the basis of βAR binding/PKA signaling, therefore, we hypothesized that we would observe an increase in respiration related pathways similar to ISO treatment. As expected, and according to our hypothesis, all respiration-related pathways induced by ISO were also induced by mAb 2C.4, confirming that 2C.4 induces respiration, and validates our analytical approach (**Figure 5B**). We then compared pathways related to fibrosis between 2C.4 and ISO treatment (**Figure 5A**), and found that like respiration pathways, mAb 2C.4 also directly induces many of the same fibrosis pathways induced by ISO in H9c2 cells. We interrogated the GSEA plots of the seven overlapping pathways (**Figure 5B, inset**) to assess the maximum enrichment score (ES_max_), and found that ES_max_ for fibrosis pathways induced by 2C.4 were at similar or greater magnitude than that of ISO. Because βAR stimulation is known to cause inflammatory changes, (75) we sought to compare inflammatory pathway changes between both treatment conditions (**Figure 5, top panel, cluster C**). However, ISO treatment caused significant upregulation of several pathways of negative T cell regulation while mAb 2C.4 treatment did not induce any inflammatory pathways overlapping with ISO treatment (not shown). The response to mAb 2C.4 was largely composed of positive T cell activation pathways, likely due to secreted factors present in mAb/B cell hybridoma culture medium conditioned with 2C.4 clones and was therefore not investigated further. We also used the ranked gene list in this analysis as input to STRINGDB for a parallel analysis of enriched pathways. This analysis yielded a similar result, with mAb 2C.4 causing enrichment in fibrosis and respiration/adrenergic pathways similar to ISO treatment (**Supplementary Figure 5a-d**). Overall, our data shows that the myocarditis-derived cross-reactive CM/βAR mAb 2C.4 induces a transcriptional response consistent with a fibrosis signature and suggests that CM/βAR AAbs in myocarditis and progressive heart failure may contribute to cardiac fibrosis.

### CM IgG autoantibodies were detected in biopsies from patients with myocarditis

EAM studies in our laboratory previously demonstrated that CM AAbs targeted the heart cell surface, and the heart as evidenced by in vivo IgG myocardial deposition and AAb passive transfer of DCM (14). To assess cardiomyocyte binding of antibodies in human tissue, we de-paraffinized myocarditis biopsy sections and incubated them with anti-human IgG or control antibodies. Biopsy sections showed diffuse IgG deposition in the myocardium of myocarditis biopsies(n=5) while conjugate controls (n=5) and a representative healthy subject myocardium (n=1) did not show IgG deposition within heart tissue (**Supplementary Figures 6**). IgG deposition detected in this assay represents endogenous IgG bound to myocardium.

## Discussion

AAbs that target the heart in myocarditis have been previously described, yet no human studies have demonstrated a connection between CM and βAR autoantibodies (AAbs) in nonrecovery of myocarditis/DCM patients. Several novel findings in our study suggest a link between these autoantibodies and outcomes of myocarditis/DCM: 1) CM AAbs reactive to unique peptide epitopes in the S2 hinge region are significantly elevated in the group of myocarditis patients that fail to recover LVEF and develop progressive heart disease. It is important to emphasize that CM AAbs in nonrecovered patients are elevated significantly above those in myocarditis patients who spontaneously recover LVEF and may therefore be prognostic of nonrecovery; 2) CM AAbs may be functional biomarkers that induce gene expression consistent with known features of pathogenesis, namely fibrosis and apoptosis; 3) Endomyocardial biopsy from myocarditis patients with positive IgG deposits frequently demonstrated a high degree of fibrosis.

Although we report here specifically on AAbs as a potential pathogenic mechanism leading to fibrosis and poor outcomes, both T cells and macrophages as well as other immune mediators are also involved (39,76) as shown in our model diagram in Figure 6. Th17 cells are important in driving IgG production as well as producing IL17A and responding to elevated IL-6 elevated due to inflammation or infection. IL6 and IL17 were both biomarkers affected in nonrecovery and Th17 cells clearly associated with worsening heart failure as measured by the NYHA(39). And further, dendritic cells respond to environmental cues where IL10 is protective and IL17A/γIFN is inflammatory and destructive in Lewis rat EAM (77).

**Figure 6:**
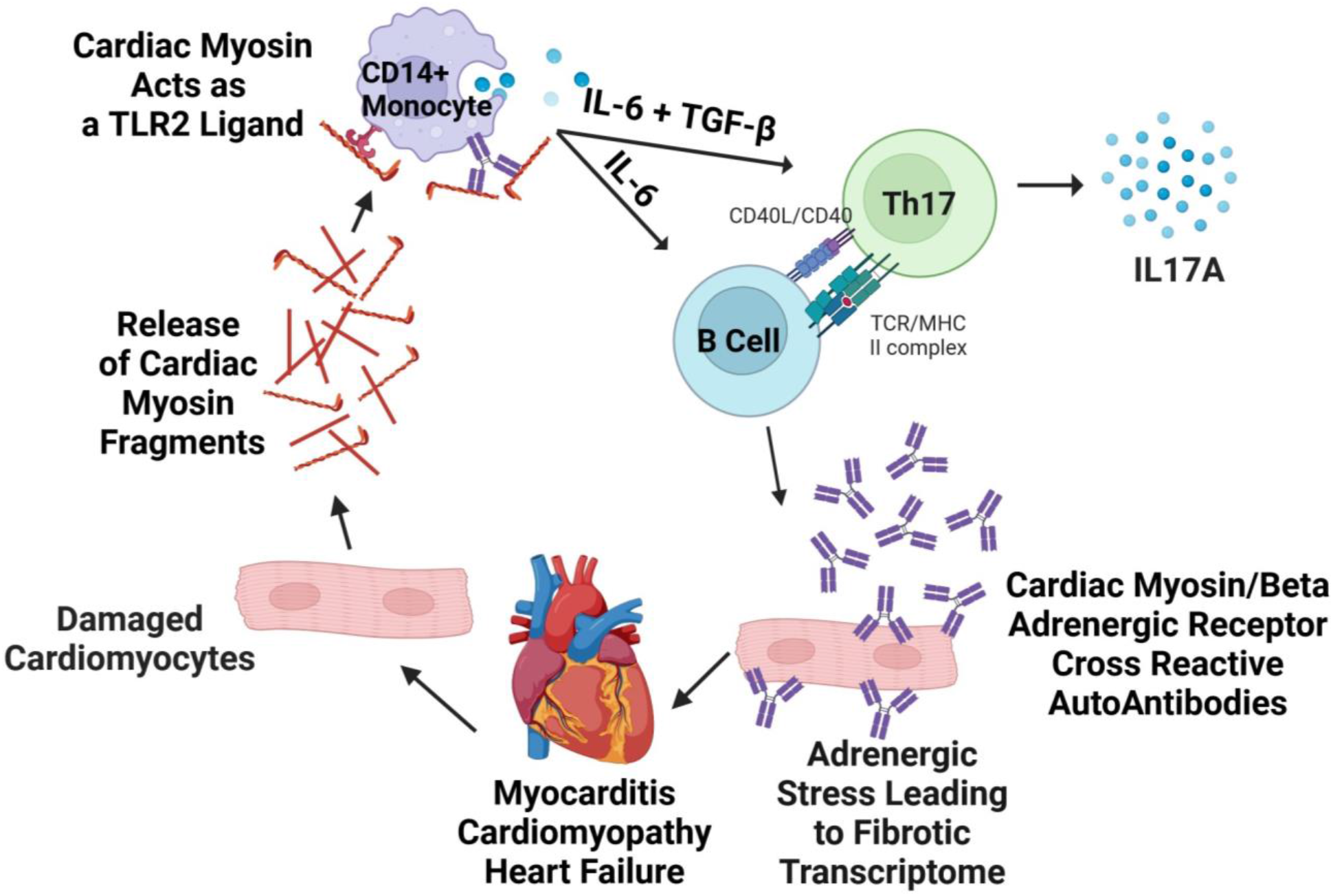
Model of pathogenic mechanisms in human autoimmune myocarditis. Lytic viral infection of cardiomyocytes or other causes of cell damage releases cardiac myosin (CM) which acts as a danger signal via TLR2/8, causing cytokine secretion that can be suppressed with TLR2 blocking antibody *in vitro*. In human myocarditis, cytokine secretion responses from CD14^+^ monocytes are exaggerated in comparison to healthy and promote Th17 differentiation. Antibodies to CM are also produced, and agonize βARs on cardiomyocytes, directly inducing a pro-fibrotic response in cardiomyocytes and potentially other cell types in the myocardium. Created with BioRender.com and modified from Myers 2016 ^6^.

The potential role of CM AAbs in poor outcomes is supported in several other studies, including one of a myocarditis cohort (78), and another in cardiac events in type 1 diabetes (47). Similarly, another group recently demonstrated that intracellular CM binding protein-C AAb positivity was prognostic for poor outcomes in patients with acute coronary syndrome (79). Other evidence shows that CM AAbs arise in settings where cardiomyocyte death occurs and even in healthy subjects, albeit at low frequencies (80). In myocarditis, patients have been reported to have CM AAbs at a rate of 17 (81) to 52 percent (82). Patients with dilated cardiomyopathy have been found to have CM AAbs at a rate of 20 (83) to 27 percent (84). Our findings also parallel the data presented by Simpson and Cantor, et al. who found significantly higher CM AAbs in baseline serum samples from newly diagnosed groups of pediatric myocarditis/DCM patients compared to healthy subjects, and higher PKA activation at the early time point in those patients who did not recover later in disease (85). Just recently, the hypothesis that the βAR contributes to poor outcomes was tested in a model of PD-1 related myocarditis using repeated adrenergic stress-induced myocarditis in PD-1 deficient mice (86).

The potential clinical significance of AAbs to the β1-adrenergic receptor (β1AR) have been recognized with the finding that the second extracellular loop has B and T cell epitopes (87) and that AAbs to this domain have potential agonistic activity in myocarditis and other cardiovascular diseases (88). Several other studies of non-specific immunoadsorption demonstrated that IgG removal improved cardiac function which coincided with reduction in β1AR AAb titers (89,90) and cardiac improvements persisted to one year after immunoadsorption treatment (91). Patients with β1AR AAbs benefit from both non-specific and β1-specific immunoadsorption (92); however, multiple studies have shown that β1AR AAbs reappear in some patients, coinciding with worsening of cardiac function (92,93). Similarly, passive serum transfer studies demonstrated that β1AR AAbs have the potential to induce cardiomyopathy in the absence of other infectious or immune challenge (18). Physiologic β1AR signaling in the heart via catecholamines promotes contractility and heart rate, but overstimulation induces apoptosis (14,94) and heart failure in murine models (95,96). Previous work in our laboratory used the Lewis rat model of EAM to demonstrate AAbs develop against the CM molecule and concomitantly induce PKA signaling in cardiomyocytes (14). Other studies, including work from Caforio et al. (84) and de Leeuw et al. (97) suggest that cardiac myosin AAbs are early markers of idiopathic cardiomyopathy and heart-specific AAbs diminish late in disease progression. CM AAbs have also been associated with rheumatic heart disease and autoimmune responses against group A streptococci (98). These responses were also cross-reactive with the coxsackievirus suggesting that infection could be an eliciting agent of these AAbs in some instances.

Our previous work on CM autoantibodies from human myocarditis led us to hypothesize that they may contribute to poor outcomes and may be useful for prognosis. Initially we tested this by measuring myocarditis patient autoantibodies reactive to whole cardiac myosin (CM) isolated from human heart tissue. In our cohort, myocarditis patients had significantly elevated CM AAbs relative to healthy subjects at baseline (**Fig. 1)** AAbs to whole CM did not correlate with EF in patients who recovered (**Supplementary Fig. 1**) but did correlate with EF in nonrecovered patients with poor EF outcomes at 1 year (**Fig 1d**.). To confirm the CM reactivity by these AAbs, we then tested sera from our cohort using 25-mer synthesized peptides from the most immunogenic region (S2 fragment) of the human CM molecule (98,99). Using this approach, myocarditis patients who failed to recover EF by one year had CM AAbs significantly elevated at baseline compared with patients who recover EF by one year (**Fig 1c**). Several CM S2 Peptides were elevated compared to healthy subjects (**Supplementary Fig. 1**). Nonrecovered patients had unique reactivity to peptides 1, 17, 29, and 30, which were significantly elevated in our present cohort. We observe notable similarity in peptide reactivity in a previous cross-sectional study using the same 25-mer peptides examining diabetic cardiomyopathy, myocarditis, and DCM patients (20). In the work by Mascaro-Blanco et al., peptides 1, 9, 17, and 30 were elevated in myocarditis or DCM, and in diabetic cardiomyopathy, peptides 6, 8, 10 and 28 were elevated. Sequence alignment between the S2 25-mers used in our assay and β1AR and B2AR revealed that a locus of CM corresponding to the S2-3 and S2-4 peptides (MYH7 sequence VQAEQDNL) had a 62.5% overlap with a section of the βAR2 in the third extracellular loop (Supplementary Table 3). Similarly, peptide S2-30 (MYH7 sequence RSKAEETQRSVND) shared 6/13 (43%) homology with the sequence RAESDEARRCYND occurring in the β1AR second extracellular (EC) loop (Supplementary Table 3). This locus of the β1AR, when immunized in Wistar rats, produced an autoimmune response specific to only the β1AR and created a positive chronotropic effect (100). Additionally, autoantibodies to the second EC loop of the β1AR predicted death of patients with idiopathic DCM (101). Based on these observations, we suggest that in settings of cardiac damage, antigenicity to βARs may occur frequently at several regions across the CM molecule (Supplementary Table 3).

Our previous work on CM autoantibodies led us to hypothesize that due to their functional cross-reactivity with the βARs, myocarditis patient outcomes could be predicted by both serum CM AAb concentration and PKA activation. While both recovered and nonrecovered patients had elevated PKA activity and CM AAb concentrations compared with healthy subjects, PKA activity was not directly and statistically associated with nonrecovery and poor outcomes in the same way the anti-cardiac myosin AAbs were **(Figure 3b).** This was surprising given that antibodies specific for cardiac myosin have been established to cross-react with βARs and induce PKA signaling (14).

However, AAb interactions with GPCRs goes beyond direct agonization, and includes allosteric inhibition or activation of the endogenous ligand, alteration of Gα subunits, and other mechanisms outside the scope of this discussion but reviewed recently (102). Along those lines, AAbs to β1AR have also been shown to sensitize the β1AR to agonization by isoproterenol by a factor of 3 (103), therefore it is reasonable to consider that poor outcomes are a result of modulation of the β1AR signaling that were not assessed in the present study. Another potential contribution of CM AAbs to poor outcomes is that they have cross-reactivity with other targets. We used basic local alignment search tool (BLAST) to screen for similar sequences to the S2 CM peptides used in our assay to generate hypothetical cross-reactive targets. Results were considered potential targets if they had an extracellular domain, had 5 or more homologous amino acid residues, were expressed in relevant heart cell types (https://www.proteinatlas.org) (104), and lastly if the homologous amino acid sequence was a predicted B cell epitope (105). These predicted B cell epitopes are presented in **Supplementary Table 4**.

Our previous studies on the immunogenic properties of CM provide other mechanistic insight of CM-AAbs and potentially extracellular CM bound to CM-AAbs in immune complexes. CM liberated due to cardiomyocyte cell death functions as a damage-associated molecular pattern (DAMP) that signals via Toll-like receptor 2 (TLR2) and TLR8, eliciting elevated Th17-promoting cytokine secretion in monocytes (106) **(Figure 6)**. Notably, CD14+ monocytes exposed to CM in vitro secrete Th17-differentiating cytokines IL-6, TGF-β, as well as IL-23, an effect observed in healthy subjects and significantly more in CD14+ monocytes from myocarditis/DCM patients (6) (**Figure 6**). TLR2-blocking antibody prevented CM induced cytokine secretion in myocarditis patients-derived monocytes, confirming that in humans, CM signals via TLR2 on CD14+ monocytes. Importantly, our previous study found that TGFβ was significantly elevated in nonrecovered myocarditis/DCM patients(6). Beyond playing a role in Th17 differentiation, TGFβ is notorious in fibrosis induction(107,108), and a major contributor to the pathogenesis of DCM(109,110).

Our analysis of transcriptomic changes in H9c2 cells after stimulation with myocarditis patient-derived human mAb 2C.4 showed a marked increase in fibrosis-associated pathways (**Figure 5**). Established in 1976 (111) H9c2 cells are a common model of cardiomyocytes despite having no spontaneous beat (112), and are used in hundreds of cardiovascular studies each year, especially in toxicology studies. We chose H9c2 cell line anticipating an acute toxic effect of mAb 2C.4 due to overstimulation of βARs, and the observation of upregulated fibrosis pathways was an unexpected finding. H9c2 cells are also a useful model for fibrosis given that a proteomic analysis demonstrated that they produce Col1a1, Col1a2, and Col3a1 (113), which are ordinarily found in fibroblasts (114). Our previous work demonstrated that CM AAbs cross-react with βARs, so it is a reasonable expectation that in patients with elevated CM AAbs, βAR signaling would affect a wide variety of cell types. β2AR signaling on fibroblasts was shown to induce proliferation and production of IL-6 (115), and our analysis of H9c2 cells demonstrates an upregulation of fibrosis transcripts, therefore it is reasonable to speculate that because CM AAbs have affinity to β2AR (**Supplementary Figure 3**), they may exacerbate progressive fibrosis by inducing extracellular matrix secretion, proliferation of fibroblasts, or stimulation of collagen production in myofibroblasts. The contribution of CM AAbs to pathogenesis is likely to be variable across individual patients and individual AAbs from various B cell clones, as we have shown previously in mouse studies where high cytotoxicity correlates with higher AAb avidity (116). In absence of direct cytotoxicity, CM AAbs can likely cause other widespread effects across a variety of cell types expressing βARs (20,117) (**Figure 6**). Our previous work demonstrated that CM Aabs cross-react with βARs, so it is a reasonable expectation that in patients with elevated CM AAbs, βAR signaling would affect a wide variety of cell types. β2AR signaling on fibroblasts was shown to induce proliferation and production of IL-6 (115) and our analysis on H9c2 cells demonstrates an upregulation of fibrosis transcripts, therefore it is reasonable to speculate that because CM AAbs have affinity to β2AR (**Supplementary Figure 3**), they may exacerbate progressive fibrosis by inducing extracellular matrix secretion, proliferation of fibroblasts, or stimulation of collagen production in myofibroblasts. The contribution of CM AAbs to pathogenesis is likely to be variable across individual patients and individual clones, as we have shown previously in mouse studies that high cytotoxicity relates to higher avidity (116). It is therefore likely that CM AAbs can have other widespread effects across a variety of cell types expressing βARs.

Our present study supports the hypothesis that CM AAbs elicit direct and deleterious effects on cardiomyocytes. The transcriptomic changes induced in H9c2 cardiomyocytes by mAb 2C.4 within one hour of treatment overlapped with that of isoproterenol (ISO) inducing transcripts in cellular respiration and fibrosis-related pathways (**Figure 6**). β1AR signaling is well known to increase cellular respiration in multiple cell types (118) and has been previously demonstrated to contribute to fibrosis (119). To our knowledge, our report is the first direct demonstration of profibrotic potential from a myocarditis/DCM patient-derived mAb.

While our previously published works demonstrated that CM AAbs cross-react with and agonize βARs leading to PKA signaling (14,20), we sought to reproduce this finding in our longitudinal cohort where PKA signaling was evident throughout the year following onset of myocarditis/DCM. PKA activation was inhibited by anti-IgG beads and not BSA coated beads (**Supplementary Figure 4**), demonstrating that PKA signaling due to serum components was caused by IgG, and not by norepinephrine or other molecules. This demonstrates the importance of AAbs as β-adrenergic stressors in myocarditis and may predispose certain individuals to more protracted disease.

To further investigate the potential of myocarditis/DCM serum compared to the human myocarditis derived human mAb to induce cellular gene effects on H9c2 cells, transcriptomic analysis revealed differential expression of genes associated with heart failure, including apoptosis, fibrosis, and hypoxia pathways. To confirm the transcriptomic analysis and cellular effects, we performed a parallel experiment with a larger number of myocarditis (n=6) and healthy (n=4) sera to determine by PCR a subset of differentially expressed genes (**Figure 4E**). Several genes associated with heart failure, fibrosis and hypertrophy were differentially upregulated across both experiments: XDH, NFATC2, CAMTA2, and ITGB6. We have previously found that myocarditis patients frequently have elevated IL-6, suggesting the possibility of NFATc2 gene expression is induced by IL-6 receptor engagement, which is known to induce cardiomyocyte hypertrophy in vitro (120–122). Similarly, we found elevated expression of CAMTA2, a known positive regulator of cardiomyocyte hypertrophy and proliferation, and increased cardiac hypertrophy in transgenic mice overexpressing CAMTA2 (123). Integrin αvβ6 (ITBG6) binds TGFβ1 and may be involved in activating fibrosis in the heart. Direct binding of integrin αvβ6 to the latency-associated peptide of TGFβ was demonstrated by Munger et al (124). In their model of pulmonary fibrosis, mice lacking αvβ6 were protected from fibrosis. It was later shown by Shi et al. that activation of TGFβ by αvβ6 did not occur by binding alone, but additionally required the transmission of contractile force via αvβ6 (125), typically thought to be exerted by myofibroblasts in the setting of cardiac injury. While speculative, upregulation of αvβ6 by fibroblasts or cardiomyocytes in response to CM AAb suggests another pathway contributing to cardiac fibrosis in myocarditis/DCM patients.

When we compared our gene expression results from patient sera with that of H9c2 cardiomyocytes we found that most DE genes (238 of 259) induced by myocarditis/DCM patient sera were also DE by mAb 2C.4 and had the same direction of change. The fact that AAbs in patient sera elicited a similar effect to human myocarditis-derived mAb 2C.4 implies our study has relevance to human disease. We hypothesize that CM AAbs contribute to pathogenesis through their cross-reactivity to βARs, where βARs signaling is pleiotropic, including cytotoxic and fibrotic, and otherwise physiologically altering cardiomyocytes (14,126), or causing arrhythmias (127,128), thereby contributing to poor outcomes. Previous studies have suggested that anti-β1AR AAbs may lead to poor outcomes including myocardial scarring, cardiomyocyte toxicity, and heart failure due to excessive stimulation of the β-adrenergic receptor (18,129). When complexed with extracellular cardiac myosin, CM AAbs may deposit in the heart or circulate and activate macrophages and monocytes through TLRs (106) and/or through FcRs. Other lines of evidence support this hypothesis such as mice lacking a functional TLR2 gene had diminished cardiac fibrosis after myocardial infarction (130,131). To date, no study has demonstrated a connection in human myocarditis between highly elevated CM AAb binding directly in the myocardium with direct affects where the beta receptor on heart cells leads to an apoptosis and fibrosis signature causing heart injury and poor outcomes in the nonrecovered myocarditis/DCM group. Although our present study has not directly measured apoptosis or fibrosis, we have provided data suggesting a causal relationship.

The major limitation of our study is the small number of human samples studied which is often the case in rare diseases like myocarditis. However, importantly, there are few studies that have investigated a longitudinal group of humans with myocarditis/DCM where recovered and nonrecovered groups could be studied with a focus on biomarkers associated with poor outcomes. Although studies by Caforio et al and Schultheiss et al are the most notable that support our findings (11,132), no studies have connected the pathogenic relationship of CM, βAR, fibrosis and cardiac recovery as we have in this study. Validation of elevated CM AAbs as markers of LVEF recovery status is justified in a larger cohort study of myocarditis/DM, and if validated they and other inflammatory biomarkers may become useful tools in identifying high risk of nonrecovery in myocarditis/DCM in clinical practice.

In conclusion, the major new finding in our study is the demonstration of CM AAbs as a potential biomarker of poor outcomes in myocarditis. Secondly, using a novel myocarditis-derived human CM mAb, we provided evidence that CM AAbs are capable of directly inducing fibrosis gene pathways and may therefore contribute to cardiac fibrosis in myocarditis or other cardiomyopathies. Further investigation of CM AAbs may lead to their establishment as a useful prognostic for nonrecovery of LVEF. In summary, our study provides important new insights into autoimmune mechanisms in human myocarditis/DCM suggesting CM AAbs are a biomarker to identify risk of nonrecovery in patients with myocarditis.

## Acknowledgments

The authors would like to thank Damita Jo Carryer for patient recruitment and Kathy Alvarez for her expert technical assistance. We thank Dr. Stanley Kosanke, pathologist at the OUHSC, for his assistance with the human myocarditis biopsy specimens. Funding included an American Heart Association predoctoral fellowship award to CS, the National Heart Lung and Blood Institute R01HL56267 and R01HL135165 to MWC and LT, the National Institute of Allergy and Immunology T32AI00763 (MWC) support for CS and JM, and the Myocarditis Foundation fellowship award to JM. MWC was the recipient of an NHLBI MERIT Award. Dr. Fairweather was supported by NIH R21 AI152318, R21 AI145356, R21 AI154927, R01 HL164520, and American Heart Association 20TPA35490415, as well as institutional support from Mayo Clinic Center for Regenerative Medicine in Florida. Dr. Katelyn Bruno was supported by NIH R21 AI163302, R21 AI180863-01A1, American Heart Association award number 23SCEFIA1153413, and Department of Defense CDMRP PR210385.

